# Localization of multiple hydrogels with MultiCUBE platform spatially guides 3D tissue morphogenesis *in vitro*

**DOI:** 10.1101/2022.09.08.507064

**Authors:** Kasinan Suthiwanich, Masaya Hagiwara

## Abstract

Localization of multiple hydrogels is expected to develop the structure of 3D tissue models in a location specific manner. Here, we successfully localize morphogenesis within individual tissues by exposing different hydrogel conditions to different parts of the tissues. We develop a unit-based scaffold with a unique frame design to trap hydrogel solutions inside their designated units. Interestingly, this unit-based scaffold within an optimal range of dimensional size and surface wettability can trap several cubic millimeters of hydrogels. This localization capability enables the spatial organization of hydrogel compositions, growth factors and physical conditions, as well as the position of biological samples (cells, spheroids, reconstituted tissues) relative to each hydrogel compartment. We succeed to localize the branching development of reconstituted human epithelial tissues according to the localized biomolecular and physical cues from hydrogels, regardless of the initial tissue configurations. Unlike 3D-bioprinting or microfluidics, the localization with this unit-based scaffold requires only manual pipetting and handling without any specialized equipment or skills, thus ready to use by researchers from any field. This scaffold-based localization provides a new promising route to spatially control morphogenesis, differentiation, and other developmental processes within organoids or other 3D tissues, resulting in 3D functional models for practical biomedical applications.

## Introduction

During prenatal development, human cells undergo extensive structural development into various tissues and organs with sophisticated, yet specific, architectures. For instance, a tubular bud of endodermal cells can grow and branch into the highly specific lung structure through various cell-cell and cell-matrix interactions^1^. Such physiological development has inspired researchers to recapitulate the structure and function of human organs through three-dimensional (3D) tissue modeling. Indeed, many breakthroughs over the past decade came from successful differentiations of various organ-specific, multicellular tissues called organoids^2,3^. Nonetheless, despite their consistent cellular profiles, many of these 3D tissue models have random unspecific structures, mostly in a globular shape^4,5^

This structural immaturity largely arises from the uniformity of *in vitro* conditions where cells are commonly embedded in a homogeneous hydrogel under a culture-medium submersion. Conceptually, this uniform condition exposes cells to nearly the same biomolecular and mechanical cues within the hydrogel from all directions, leading to a similar growth rate and isotropic development into a globular lump of tissue. *In vivo*, however, different parts of an embryo experience different proteins and macromolecules from the surrounding extracellular matrices (ECMs)^6^. This spatial organization (*i.e.,* localization) of ECMs guides local cells at different tissue parts to grow at different rates and develop different patterns into various location-specific structures (*i.e.*, morphogenesis). A similar dependence on ECM localization also occurs in adults, particularly during tumor/cancer development and wound healing^7,8^ Because hydrogels serve as ECM substitutes in 3D tissue modeling, the localization of different hydrogel conditions is expected to guide the morphogenesis at different tissue parts into desirable location-specific structures *in vitro*.

Although several existing platforms can localize hydrogels into multiple-hydrogel systems, their technical drawbacks limit the application of hydrogel localization in 3D tissue modeling. Microfluidic chips can fabricate both discrete hydrogel compartments and continuous hydrogel gradients by flowing different hydrogel solutions through microscaled channels^9^. However, this microfluidic method depends on optimal matching between hydrogel rheological properties and specific channel designs, thus requiring engineering skills in fluid mechanics and access to lithographic facilities. On the other hand, bioprinters can fabricate free-form multiple hydrogels through either extrusion or stereolithographic methods^10,11^. However, almost all bioprintable solutions (*i.e.*, bioinks) are molecularly functionalized or supplemented with artificial additives to gelate and stabilize the printed hydrogels against structural collapse^12^. These chemical modifications certainly affect cellular responses, which call for extensive confirmations on the suitability of a specific bioink to a specific cell type. Altogether, the current practice of hydrogel localization requires critical considerations on hydrogel properties and fabrication process. Moreover, the integrations of either microfluidics and bioprinting to 3D *in vitro* culture usually require cells or biological samples to be resuspended in hydrogel solutions prior fabrication, thus exposing cells to undesirable stress during fabrication (*e.g.,* high pressure, UV irradiation, free-air exposure)^13–15^. Consequently, the application of multiple-hydrogel localization in 3D tissue modeling to control tissue morphogenesis is very rare and requires substantial interdisciplinary expertise.

Moreover, applying multiple-hydrogel localization in 3D tissue modeling should satisfy the following end-point demands. Firstly, many biological researchers prefer hydrogels from ECM protein (*e.g.,* collagen, basement membrane extract, fibrin). These hydrogels generally require >10 min incubation at 37°C to complete gelation^16,17^, which usually form soft structures. Thus, the stabilization and localization of these ECM-protein hydrogels without any chemical modification is needed. Secondly, many researchers have already established their 3D tissue models in various shapes and sizes, from a microscaled spheroid to a tissue fiber of several millimeters (mm)^18^. Preferably, multiple-hydrogel localization should be adaptable to these existing models without complicating the already established protocols. Thirdly, the exposure to culture media critically affects the response of 3D tissue models. To distinguish the effect of hydrogel localization from the effect of medium directionality, 3D tissue models should experience a culture medium from all directions. Therefore, the multiple-hydrogel localization for 3D tissue modeling should be 1) applicable to any hydrogel including ECM-protein hydrogels; 2) customizable to various forms and protocols of established 3D tissue models; 3) devoid of any materials modification or uncontrolled fabrication stress which could alter cellular responses; 4) permeable to culture media from all direction; and 5) easy to perform with no need of sophisticated skills or tools.

Addressing these outlined limitations and demands requires a new platform to localize multiple hydrogels for 3D tissue modeling. The platform should completely localize ECM-protein hydrogels in their liquid phase *prior* incubation to allow subsequent embedding and seamless connections among hydrogel compartments. This requirement parallels concurrent developments of lattice/mesh structures to stabilize liquid thin-films by their surface tension and guide a solution flow through the capillary action^19–22^. In 3D tissue modeling, ECM-protein solutions are either placed on a solid substrate to form a hydrogel dome (**Fig.1A**) or filled in a container (*e.g.*, PDMS mold, well plate) to form a templated shape. However, these platforms failed to maintain the shape and partition of localized solutions (**Fig.1B**), in addition to their uneven medium exposure. We instead developed a unit-based scaffold with a unique cubic unit design to trap hydrogel solutions in their designate units (**Fig.1C**), which could be customized into many shapes and arrangements to allow various hydrogel configurations. Here, we established this cubic unit-based scaffold as the new platform for multiple-hydrogel localization, called MultiCUBE (**Fig.1D**). MultiCUBE successfully localized different conditions of ECM-protein hydrogels (*i.e.,* matrix compositions, presence of growth factors, crosslinking), to guide the self-organization of endothelial cells as well as the morphogenesis of epithelial tissues into specific structures at specific locations (**Fig.1E**). This platform-enabled localization of multiple hydrogels has the potential to push many existing organoids and 3D tissue models closer to recapitulating the specific structural architecture of *in vivo* physiology.

**Fig.1:**
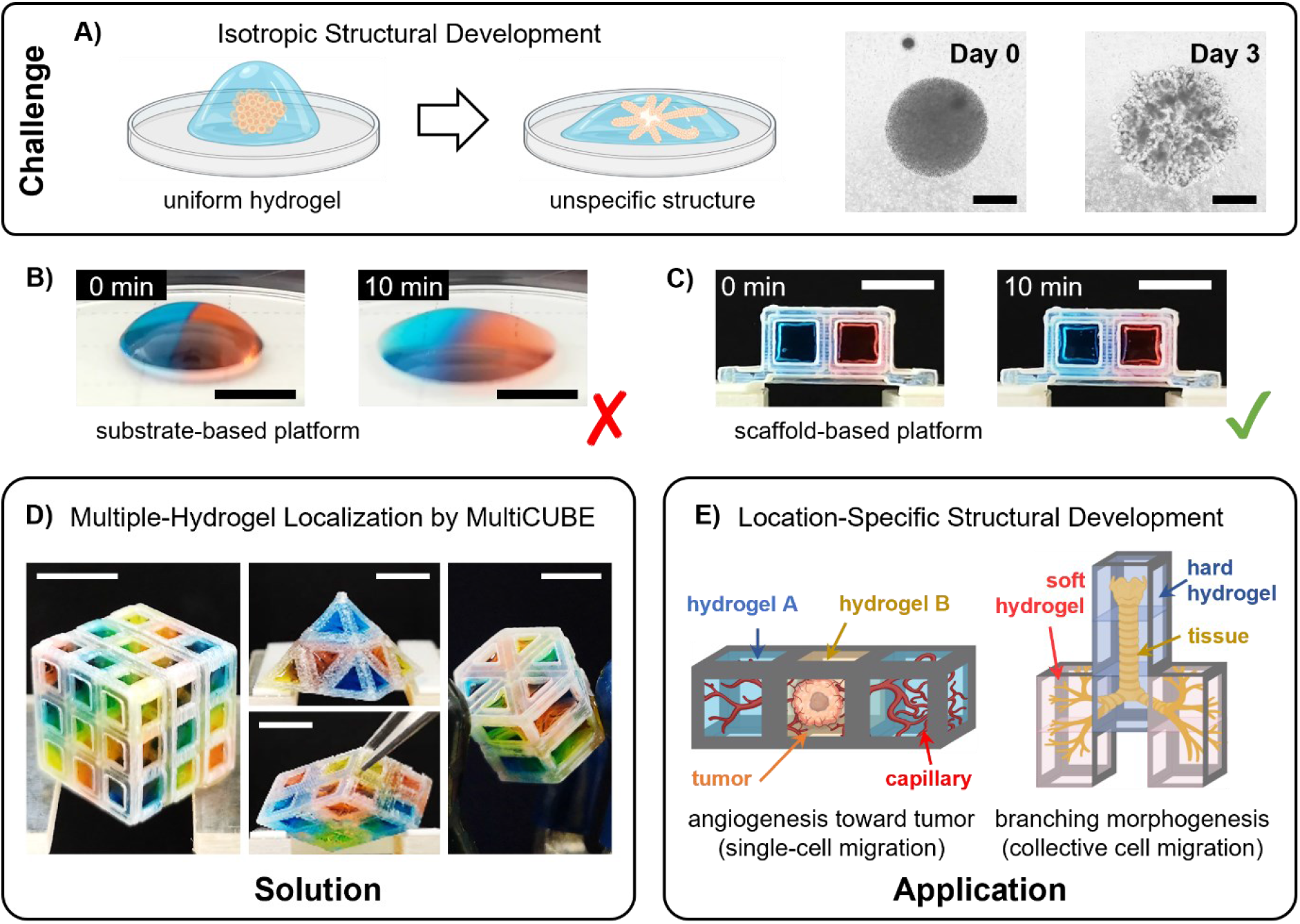
Overview of MultiCUBE development for multiple-hydrogel localization in 3D tissue modeling. **A)** Hydrogel uniformity leads to an isotropic development of 3D tissue models into unspecific structures. A high-density NHBE cluster developed into 3D tissues branching in all directions within three days in HUVEC-resuspended Matrigel dome. Scale bar: 500 μm. **B)** A hydrogel dome on a solid substrate failed to maintain the shape and partition between localized solutions. Scale bar: 5 mm. **C)** A scaffold-based platform with a specific design effectively localized two hydrogel solutions in their designated units. Scale bar: 5 mm. **D)** The resulting MultiCUBE platform could be customized with many unit shapes and arrangements for various configurations of multiple-hydrogel localization. Scale bar: 5 mm. **E)** Multiple-hydrogel localization in MultiCUBE enabled the control of location-specific structural development of various 3D tissue models based on the localized hydrogel condition in each unit.

## Methods and Experiments

### Device Fabrication

All cubic units and MultiCUBEs were 3D-printed with an acrylic resin (AR-M2, Keyence) by a commercial inkjet system (Agilista 3200, Keyence). After manually removing the supporting material (AR-S1, Keyence), all structures were cleaned by a series of solvent sonication at room temperature for 10 min each. The sonication series were i) Keyence cleaning solution twice; ii) IPA once; iii) tap water twice; iv) 70% EtOH once; and v) DI water several times. The cleaned structures were then irradiated with UV light for 10 min to completely crosslink any residual resin inside the structure.

For the solution-trapping experiment, further surface treatments were applied to the cleaned structures. Surface hydrophilization was performed by an air-plasma treatment for 3 min (PIB-10, Vacuum Device), followed by an immediate submersion under DI water for ≥10 min. Hydrophobization was done by dip-coating under a water-proof solution (Daiso) for 10 sec and air-drying. Sessile drop measurement was conducted to measure the static water-contact angle (WCA) to confirm the success of surface treatment.

For all 3D culture experiment, cleaned MultiCUBEs were further coated with 0.3% w/v Sudan Black B (SBB; Fujifilm Wako Chemicals) in 70% EtOH for 1-2 hr to reduce the autofluorescent background from the acrylic resin^23–25^. They were then sonicated with DI water and 70% EtOH alternately to remove excessive coating. Coated MultiCUBEs and other cleaned structures (*i.e.*, holders, templating jigs) were transferred into a biosafety cabinet under a submersion in 70% EtOH. After drying with aspiration, they were sealed in a well plate, sterilized with UV light for 10 min, and kept at 4°C until further use.

### Solution-Trapping Experiment

Under an ice bath, acid-hydrolyzed porcine collagen solution at (3 mg/ml; Cellmatrix Type I-A, Nitta Gelatin) was neutralized with the reconstitution buffer solution (dyed with food coloring) according to the manufacturer instruction and used immediately. All solutions were kept in an ice bath throughout the experiment. The volume pipetted into each test cube corresponded to S^3^ μl (or mm^3^) with 5-10 μl surplus to compensate for the viscosity-induced inconsistency during pipetting. Images were taken and analyzed with ImageJ software.

### Cell Culture

Normal human bronchial epithelial cells (passage 4-6, NHBE; CC-2541, Lonza) and human umbilical vascular endothelial cells (passage 4-6 HUVEC; C2519A, Lonza) were cultured in BEGM™ BulletKit™ (CC-3170, Lonza) and EGM-2 BulletKit™ (CC-3162, Lonza), respectively, for two days before 3D culture experiment. Mouse pancreatic islet endothelial cell line (passage 8, MS1; ATCC), human hepatocellular carcinoma cell line (passage 4, HepG2; ATCC), and normal human lung fibroblasts (passage 4, NHLF; CC-2512, Lonza) were cultured in DMEM +10% FBS (Thermo Fisher). All media were supplemented with 50 IU/ml penicillin and 50 μg/ml streptomycin (Thermo Fisher).

### 3D Culture Experiment: Tumor/Angiogenesis Model

The following method was modified from a previously established protocol^26^. HepG2 and NHLF were resuspended in EGM-2 (supplemented with 28.2 μg/ml neutralized collagen) at the final concentration of 6.20×10^5^ and 2.81×10^5^ cell/ml respectively. 150 μl cell resuspension was filled into an agarose-coated, 96-well plate and cultured for two days to form a spheroid. On the day of the experiment, a high-density MS-1 pellet (8.15×10^4^ cell/μl) was resuspended in 1:1 Matrigel:collagen solution at the cell:solution volume ratio of 1:200 in an ice bath. Then, a pre-fibrin solution containing 107.4 μl of fibrinogen (8 mg/ml; Sigma), 18 μl of collagen (3 mg/ml; Nitta Gelatin), 3.6 μl of aprotinin (25 μg/ml; Roche) was prepared in an ice bath. The pre-fibrin solution was supplemented with 1μl of thrombin (50 U/ml; Nacalai Tesque), and immediately pipetted into the 2^nd^ unit of a 3×1 MultiCUBE. After placing a spheroid into the fibrin solution, MS1-Matrigel-collagen mixture solution was then filled into the 1^st^ and 3^rd^ units.

### 3D Culture Experiment: Tissue Branching-Pattern Model

Frozen basement membrane extract (Matrigel; 356231, Corning) was thawed at 4°C one day before use. For crosslinking-condition localization, genipin (G4796, Sigma) was premixed to Matrigel at the final concentration of 0.5 mM under an ice bath. For HBEGF localization, human recombinant heparin-binding EGF-like growth factor (HBEGF; E4643, Sigma) was premixed to Matrigel at the final concentration of 2.5 nM under an ice bath. An aluminum wire (Ø 0.3 mm, fixed on the lid of a 12-well plate) was coated with 5% v/v LIPIDURE^®^ in EtOH (CM5206, NOF Corporation) for 1 min and left dry for at least 30 min before use. This phosphorylcholine-based coating formed bioinert nanoscaled thin-film^27–30^, which prevented mechanical damages and bubble formation inside the templated space. After trypsinization and centrifugation, HUVEC pellet was resuspended distributively in the prepared Matrigel at the ratio 1:200. HUVEC-distributed Matrigel were pipetted into each unit of 3×1 MultiCUBE, which was then placed vertically to a jig holder in a 12-well plate such that the 1^st^ unit was at the topmost position. After carefully closing the lid to insert the coated aluminum-wire mold through the centers of all three units, the plate was incubated for at least 30 min to cure and mold the localized hydrogel. Then, 7 μl NHBE pellets were seeded into the templated tubular space dropwise with a Ø 0.3 mm gel-loading tip. After letting the pellet settle down in an incubator for 20-25 min, 2.5 μl of hydrogel corresponding to the 1^st^ unit was dropped on the tube inlet and the MultiCUBE in a horizontal orientation was incubated for 10 more min. Then, the MultiCUBE was submerged in a culture medium into a 24 well-plate such that all units were under the meidum (approx. 1.75 ml). All samples were observed and imaged under a 4×, bright-field microscope daily. For all culture conditions with crosslinked-Matrigel localization and without localization, the culture medium was a 1:1 mixture of BEGM and EGM-2 (control medium). For all culture conditions with HBEGF(+)-Matrigel localization, the culturing medium was the control medium without EGF.

### Immunofluorescence Staining

The samples together with hydrogels and the MultiCUBE were rinsed with DPBS for 10 min before fixation with 4% paraformaldehyde phosphate buffer solution (09154-56, Nacalai Tesque) for 20 min at room temperature. The samples were then washed with DPBS three times, 10 min each, and permeabilized with 0.5% Triton-X for 1 hr at room temperature. Then, they were washed with 100 mM glycine in DPBS three times, 15 min each, to quench autofluorescence from unreacted aldehyde residue. Then the samples were blocked with IF solution (0.2% Triton-X + 0.1% BSA + 0.05% Tween20 in DPBS), supplemented with 10% goat serum for at least 1 hr.

The samples from tumor/angiogenesis models were stained with the primary antibody rabbit anti-CD31 (1:50; ab28364, Abcam) at 4°C overnight. Then, it was washed with IF solution at room temperature three times, each for at least 20 min. Then, the sample was stained with the secondary antibody goat anti-rabbit, Alexa Fluor 555 (1:200; A21429, Invitrogen) for 4 hrs at room temperature in the dark. Then, it was washed once with IF solution and stained with 600 nM DAPI in DPBS for 1 hr. Then, it was washed with DPBS three times, at least 10 min each before imaging.

After blocking, the samples from the tissue branching-pattern model were stained with 600 nM DAPI in DPBS and 50 μM phalloidin (PhenoVue Fluor 488; CP24881, PerkinElmer) together for 1 hr. Then, it was washed with DPBS three times, at least 10 min each before imaging. All imaging was performed with a BZ-X700 microscope (Keyence).

### Image Analysis

All bright-field images of the sample were stitched with ImageJ and the AdvanView software dedicated to the camera (AdvanVision). For all branching-pattern models, the tube perimeter and branch length were manually measured by AdvanView. Then, the data were exported *via* Excel files to MatLab for subsequent visualization and statistical analysis.

## Results

### The solution-trapping ability of MultiCUBE cubic units depends on the interplay between dimensional design and surface wettability

The key to multiple-hydrogel localization lies in the frame design of each cubic unit of MultiCUBE. For comparison, hydrogel solutions almost always drip out from a simple cubic unit with straight frames (**Fig.2A**: upper) because the solutions had no surface along the inner-cube space to adhere (**Fig.2B**: upper, dashed outline). Even if a solution sustained inside the cubic unit, filling another solution immediately caused solution shifting across units. However, each MultiCUBE unit was constructed from L-shaped frames which formed a surface band extended along the edge and a square window on the center of each cubic face (**Fig.2A**: lower). This surface extension basically served as a physical barrier against the weight of hydrogel solutions along the z axis and against the cohesion along the x-y plane (**Fig.2B**: lower). MultiCUBE units with L-shaped frames effectively trapped their hydrogel solutions inside and prevented solution shifting across neighboring units.

**Fig.2:**
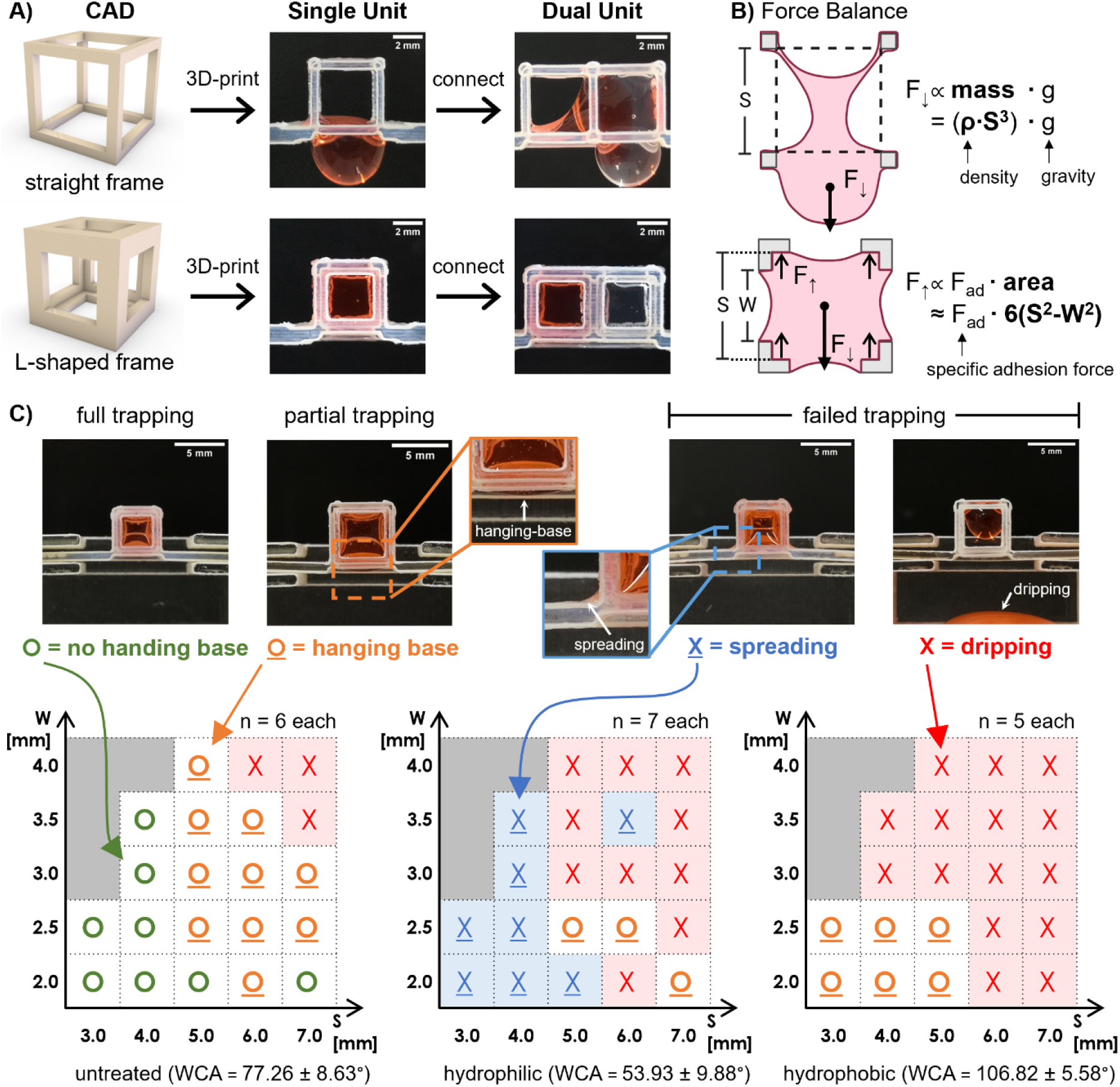
The unit design of MultiCUBE and the effect of physical parameters on solution-trapping ability. **A)** A cubic unit with straight frames failed to trap a hydrogel solution inside whereas a cubic unit with L-shaped frames successfully trap a hydrogel solution without dripping or shifting. **B)** The straight frame lacked sufficient physical barriers to counter against the hydrogel weight (F↓) whereas the L-shaped frame countered the hydrogel weight by providing surface extension along the dashed outline for solution adhesion (F↑). **C)** Different conformations of trapped hydrogel solutions indicated different solution-trapping ability - no hanging base indicated a full trapping; stable hanging base indicated a partial trapping; and spreading or dripping indicated failed trappings. These conformations varied across different inner-cube space (S) and window-frame size (W), as well as surface wettability. Each symbol in each (S,W) grid represented the occurrence of different solution conformation: **X** (dripping), **<u>X</u>** (spreading), **<u>O</u>** (hanging base), and **O** (no hanging base). If the occurrence was unrepeatable, for example, a particular grid with one dripping, two hanging-base and four spreading would be symbolized with **X**. The number of repeats in each grid is n = 6, 7 and 5 for untreated, hydrophilic, and hydrophobic frames, respectively (**Fig.S1**).

To explore the design dependence of this solution-trapping ability, a hydrogel solution was manually pipetted into each L-framed unit across different inner-cube space (S) and window-frame size (W). To mimic 3D tissue cultures with ECM-protein hydrogel, we used an ice-chilled, neutralized collagen solution due to its wider usage and better understanding of its materials properties. The result showed that the units could trap hydrogel solutions as large as S = 7 mm. As S or W increased, the trapped solution formed a convex hanging base through the open base window with an increasing height until it eventually dripped (**Fig.2C**: lower left). This gradual deterioration of trapping at higher (S,W) and high repeatability at a specific (S,W) suggested that the solution-trapping ability depended on a balance between the hydrogel weight F↓ and the surface-adhesion force F↑. Assuming a constant density and specific adhesion force, the solution-trapping ability should roughly follow the value trend of area-to-volume ratio 6(S^2^-W^2^) / S^3^ (**Fig.S1A**). However, the overall trend of the experiment results did not follow the expected value trend, pointing to more interplaying factors such as solid-liquid interaction or pipetting dynamics.

Thus, we investigated the effect of surface wettability, and observed dramatic changes in the solution-trapping ability across different (S,W). When MultiCUBE units became hydrophilic, the trapped solution tended to spread out of the inner-cube space, which occurred over a wide (S,W) range unrepeatably. Solution dripping also occurred unrepeatably over the lower S range (**Fig.2C**: lower center). These spreading and dripping across a wide (S,W) range made hydrophilic units unsuitable for multiple-hydrogel localization. On the other hand, hydrophobic MultiCUBE units could trap hydrogel solutions within a smaller range of (S,W) with hang-base conformation. Dripping almost always occurred when S ≥ 6 mm or W ≥ 2.5 mm (**Fig.2C**: lower right). The solution-trapping ability at a specific (S,W) was more repeatable in hydrophobic units than in hydrophilic units, confirming that hydrophobic units with small (S,W) were still usable for localization. This finding was particularly useful in designing MultiCUBEs for 3D culture. In this research, we needed to coat MultiCUBE with the hydrophobic Sudan Black B (SBB, **Fig.S1B**) to suppress the frame autofluorescence^23–25^, and thus we chose the design of S = 3 mm and W = 2 mm. Altogether, these results showed the flexibility of unit/frame design and material choice for fabricating MultiCUBEs with effective solution-trapping ability.

### Solution filling process determines the success of multiple-hydrogel localization

When connecting many cubic units together, the localization of hydrogel solutions dynamically depended on the filling direction and the overall unit orientation. Testing on MultiCUBEs with three consecutive units (3×1 MultiCUBE) showed that a vertical orientation of all units along the gravity vector always led to a solution dripping whenever filling the second solution from bottom up or top down (**Fig.3A**: left). A horizontal orientation also led to a localization failure whenever filling the third solution, which often shifted toward the previously trapped solution (**Fig.3A**: center). Supposedly, the increase in the overall weight and the movement of center of gravity (CG) destabilized the solution-trapping ability, especially when the CG vector pointed toward an open window. Therefore, we solved the problem by orienting all three units diagonally such that the supposed CG vector pointed toward the unit edge. Interestingly, filling the solutions from bottom up caused an immediate shifting of the second solution downward *even before* the entire 2^nd^ unit was filled. Top-down filling, however, resulted in successful localization without solution shifting (**Fig.3A**: right, **Fig.S2,3**). This observation showed that a sufficient adhesion of a subsequent solution to the frame surface critically prevented its strong cohesion to the previous solution. Therefore, we devised a body-diagonal holder to direct the CG vector toward the unit corner, which further fortified the solution stability and facilitated manual pipetting (**Fig.3B**). The body-diagonal orientation was effective in stabilizing multiple connected hydrogel solutions within their designated units.

**Fig.3:**
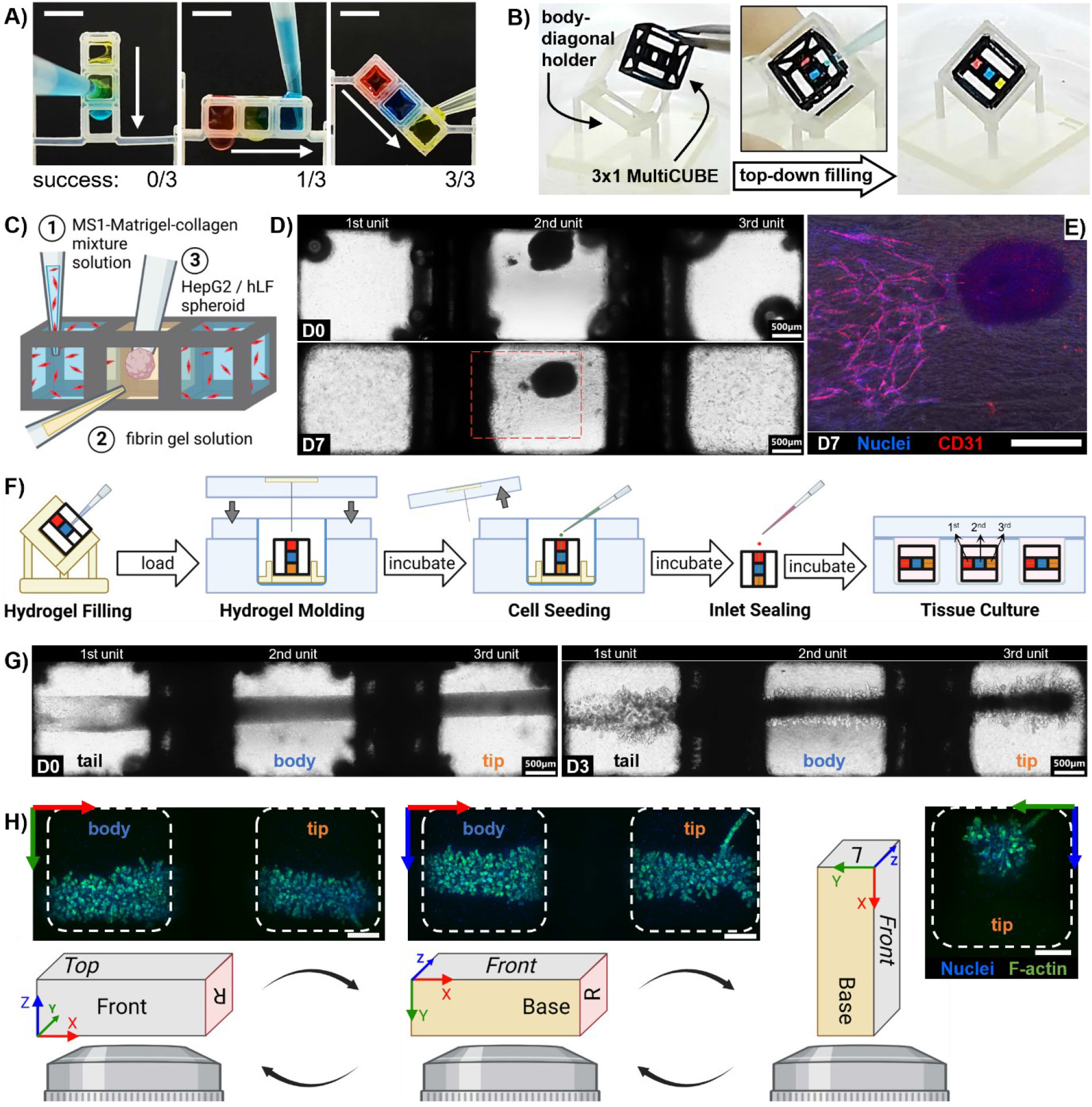
Simple fabrication process to localize multiple hydrogel solutions and embed biological samples. **A)** The diagonal orientation relative to the gravity vector led to more successes in multiple-hydrogel localization than horizontal or vertical orientations. Scale bar: 5 mm. **B)** A body-diagonal holder facilitated successful location and manual pipetting of hydrogel solutions into 3×1 MultiCUBE from top-down direction. For each unit, S=3mm and W=2mm. **C)** Biological samples such as resuspended cells or preformed spheroid could be embedded into localized hydrogel solutions. **D)** The localization of HepG2/hLF tumor spheroid in fibrin gel (2^nd^ unit) and MS1 endothelial cells in Matrigel-collagen mixture (1^st^ and 3^rd^ units) led to gradual MS1 migration toward the tumor spheroid over seven days. Scale bar: 500 μm. **E)** Immunofluorescent staining of endothelial CD31 revealed a capillary-like structure toward the tumor spheroid. Scale bar: 500 μm. **F)** The templating method could also be applied to 3×1 MultiCUBE to form a tubular space through the centers of all units for subsequent cell seeding. **G)** The resulting NHBE tubular tissues throughout all HUVEC-resuspended Matrigels underwent a branching development in a similar manner at the body and tip parts. Note that the tail part was affected by inlet sealing and its branching responses was inconsistent to the body and tip parts. **H)** The branching development on NHBE tubular tissues occurred extensively throughout its entire tubular surface, as observed by manually flipping the sample across all three axes under a fluorescent microscope.

### MultiCUBE accommodates various methods to embed biological samples into localized hydrogel solutions

Because the localization of multiple hydrogels was achieved in their liquid phase, biological samples could be easily embedded in various ways. Simply, cells can be premixed to hydrogel solutions before pipetting into each unit (**Fig.3C**: ①), or transferred as a high-density pellet or preformed spheroid into the localized solution (**Fig.3C**, ②③). We demonstrated the applicability of these embedding methods by generating a chimeric co-culture system in 3×1 MultiCUBE to mimic the endothelial angiogenesis toward a tumor^33^. A fibrin gel solution was manually pipetted into the 2^nd^ unit, and Matrigel-collagen mixture solutions with resuspended mouse endothelial cells, MS1, into the 1^st^ and 3^rd^ unit. Then, a preformed tumor spheroid of hepatocarcinoma and fibroblasts, hepG2/hLF, was transferred into the fibrin gel in the 2^nd^ unit. After curing the hydrogel and culturing under a medium submersion, endothelial cells in both units elongated, connected, and migrated across units over seven days (**Fig.3D**). Clearly, they self-organized into a capillary-like structure directing the tumor spheroid (**Fig.3E**). Similar self-organization of endothelial cells was absent when the 2^nd^ unit was filled with Matrigel instead of fibrin gel (**Fig.S4B**), highlighting the importance of appropriate localization of hydrogel compositions in modeling functional and dynamic tissue systems. This simple demonstration showed that MultiCUBE allowed the initial localization of both hydrogel conditions and biological samples, which enhanced the modeling complexity without any complicated fabrication and embedding processes.

Biological samples could also be embedded in localized hydrogels through a templating method to ensure high repeatability and consistent exposure to hydrogels in each unit^34,35^. After filling hydrogel solutions into 3×1 MultiCUBE, a thin tubular mold was inserted vertically through all three units to form a hollow space for seeding cell pellets (**Fig.3F**). We verified the applicability of this templating method by generating a tissue branching-pattern model from human primary lung cells NHBE, which was known to branch extensively in a Matrigel dome with distributed HUVEC endothelial cells within three days (**Fig.1A**)^36^. With HUVEC-distributed Matrigels in all three units, NHBE cells inside the templated space condensed into a tubular tissue within 24 hr. Then, the reconstituted tissue gradually developed branches extensively on the body and tail parts (**Fig.3G**). Note that the inlet sealing step to prevent cell leakage (**Fig.S5E,F**) disrupted the consistency of cell pellets in the 1^st^ unit, thus excluding the tail part from subsequent analyses. Observation from all three axes by manually flipping the MultiCUBE on each side revealed extensive branches covering the entire tubular surface of the tissue (**Fig.3H**), which was consistent with the branching in a Matrigel dome. Clearly, this templating method to generate a single tissue through all units could be used to investigate the location-specific morphogenesis of tissue structure.

### Localization of growth factor in hydrogel composition guided the location-specific branching morphogenesis of a single tissue

After establishing the hydrogel solution-localizing platform and the embedding method of 3D *in vitro* tissues, we focused on whether multiple-hydrogel localization could guide the morphogenesis of a single tissue into specific structures at different locations. Previously, we found that the epidermal growth factor (EGF) was the main driver of branching development in NHBE tissue through the activation of its tyrosine kinase receptor EGFR (**Fig.4A**: 1^st^, 2^nd^ column)^37,38^. However, no branching was observed on a NHBE cluster under an EGF-reduced medium when supplementing the Matrigel dome with 2.5 nM EGF (**Fig.4A**: 3^rd^ column), supposedly because EGF diffused out of Matrigel. On the other hand, supplementing Matrigel with 2.5 nM heparin-binding EGF-like growth factor (HBEGF), an EGFR-binding ligand with a higher affinity to Matrigel^37,38^, successfully generated NHBE branching under an EGF(−) medium over three days (**Fig.4A**, 4^th^ column). As expected, the localization of HBEGF(+) Matrigel only at the 3^rd^ MultiCUBE unit guided NHBE tubular tissues under the EGF(-) medium to develop branches only at the tip part (**Fig.4B,C**). We quantified the overall development by analyzing from the base-view images by considering branches with the apparent length (from the branch tip to the tube outline) longer than the apparent width (**Fig.S6**). We estimated the apparent branch density (apparent branch number divided by apparent tissue tube perimeter), which accounted for the tip contraction toward the 2^nd^ unit during the culture (**Fig.S7C,D**). We also summed up the apparent total branch length (cumulative length sum of all branches) in each tissue part to estimate the overall branching development. The results showed that in all three days of culture, the tip part in HBEGF(+) Matrigel exhibited a significantly higher apparent branch density and apparent total branch length than the body part in the control Matrigel (**Fig.4F,G**). In comparison, NHBE tubular tissues in all control Matrigel under the control medium showed no significant difference in branch formation and overall development between the body and tip parts (**Fig.3G**, **Fig.4D,E**). This comparison showed that the localization of biomolecular cues (*i.e.*, growth factor) in hydrogels successfully guided the morphogenesis of NHBE tissues into a location-specific branching structure.

**Fig.4:**
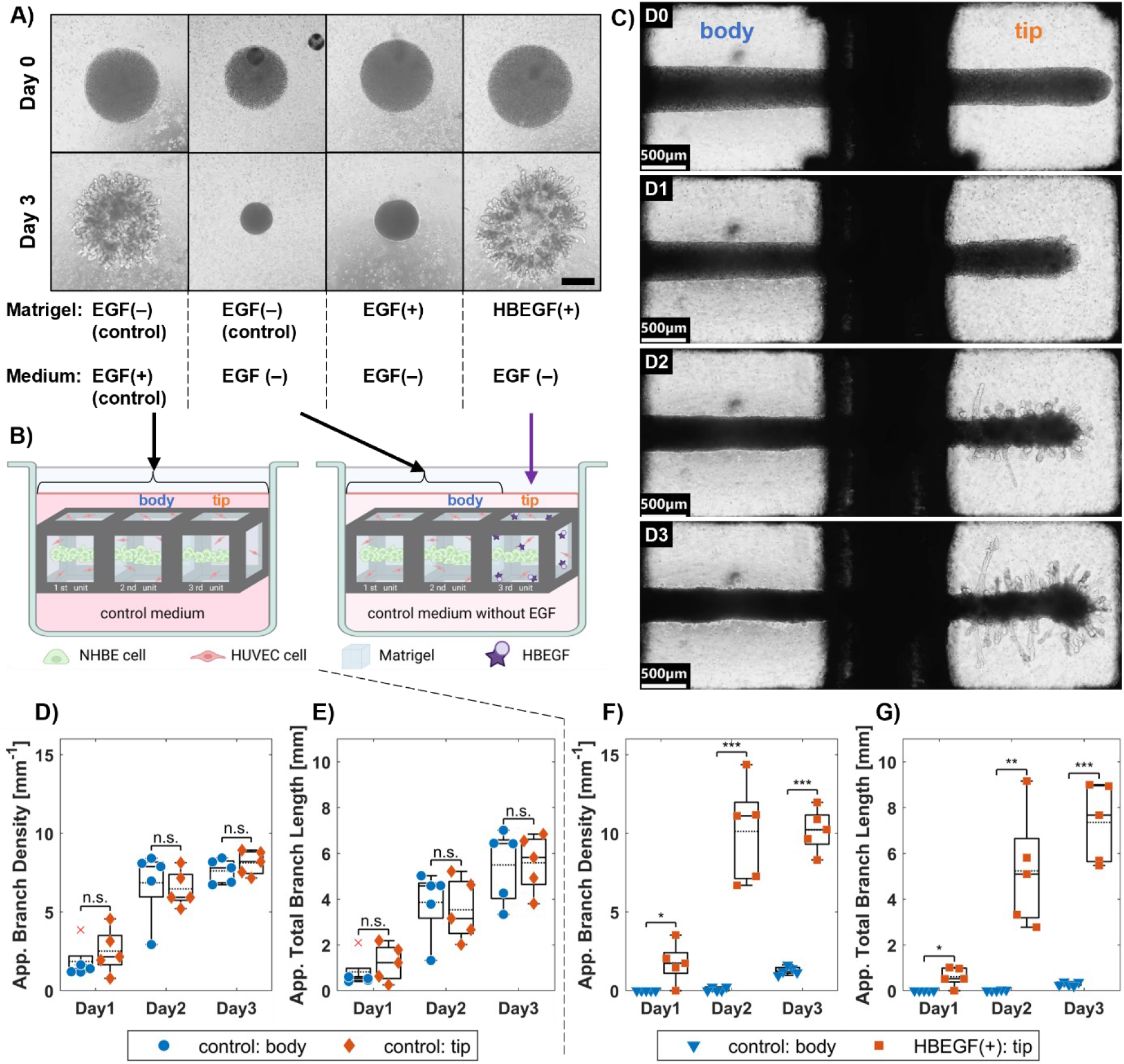
Localization of growth factor in hydrogels and the resulting branching tissue morphogenesis. **A)** EGF present in the control medium initiated the branching of NHBE tissues. Removing EGF from the control medium prevented the tissue from branching. Although supplementing 2.5nM EGF in Matrigel failed to initiate branching, 2.5nM HBEGF in Matrigel generated extensive branching from NHBE clusters. Scale bar: 500 μm. **B)** Two culture conditions were compared: the control Matrigel in all units under the control-medium submersion *vs* HBEGF(+) Matrigel localized only at the 3^rd^ unit with the control Matrigel in the 1^st^ and 2^nd^ units under the EGF(−) medium submersion. **C)** Localization of HBEGF(+) Matrigel guided the branching development only at the tip part of NHBE tissues over three days of culture. **D,E)** No significant difference in the apparent branch density and apparent total branch length was found between the body and tip parts of NHBE tissues cultured in the non-localizing condition (with the control Matrigel in all units) in all three days of culture. **F,G**) The apparent branch density and apparent total branch length in all three days were significantly higher at the tip part than at the body rd part when localizing HBEGF(+) Matrigel in the 3 unit. Data are presented in dot plot with boxplot (x: outlier, solid line: median, dash line: mean). Statistical significances are evaluated with one-way ANOVA with Tukey-Framer post-hoc test (n.s.: no significant difference, *: *p*<0.05, **: *p*<0.01, ***: *p*<0.001). Scale bar: 500 μm.

### The localized branching morphogenesis of a single tissue depends on the localized crosslinking condition of hydrogels irrespective of the initial tissue configuration

We further explored whether the localization of hydrogel physical conditions could generate a location-specific structure in a similar manner. Generally, Matrigel solution gelates into a soft gel at room temperature due to reversible noncovalent interactions among its constituent proteins^39,40^. However, the physical and mechanical conditions of Matrigel could be modified by adding genipin, a fruit-extract compound with low cellular toxicity, to form irreversible covalent bonds (*i.e.,* crosslink) among the constituent proteins (**Fig.5A**)^41,42^. This crosslinked Matrigel suppressed the branching development in NHBE tissues (**Fig.5B**), as indicated by extremely low apparent branch density and total branch length (**Fig.5C,D**). When the crosslinked Matrigel was localized only at the 3^rd^ unit, only the tip part did not develop branches along the tube surface (**Fig.5E**), showing significantly lower apparent branch density and total branch length than the body part in normal Matrigel (**Fig.5F,G**). Note that the body part of NHBE tissues swelled substantially during three days of culture (**Fig.S8G,H**). Inversely, filling the 1^st^ and 2^nd^ units with crosslinked Matrigels and the 3^rd^ unit with normal Matrigel localized the branching morphogenesis only at the tip part of NHBE tissues (**Fig.5H**). Again, the body part swelled over three days, thus showing no correlation between the swelling behavior and the hydrogel localization (**Fig.S8**). The tip part in normal Matrigel had significantly higher apparent branch density and total branch length than body part in crosslinked Matrigel (**Fig.5I,J**). Interestingly, the density range of the tip in normal Matrigel was also substantially higher and broader than the density range of the body in normal Matrigel (**Fig.5F** *vs* **Fig.5I**). This difference likely arose from more tip contraction when localizing normal Matrigel than when localizing crosslinked Matrigel in the 3^rd^ unit (**Fig.S8E,F**). Critically, we found no significant difference in the apparent branch number and total branch length when we compared different tissue parts in the same hydrogel condition (**Fig.S9**). This observation strongly suggested that the branching morphogenesis at any tissue part was determined by the absence of crosslinking, and not affected by the tip contraction nor swelling behavior. This experiment demonstrated not only the effect of localizing hydrogel physical conditions, but also the controllability of localized tissue morphogenesis regardless of the initial configuration of the tissue.

**Fig.5:**
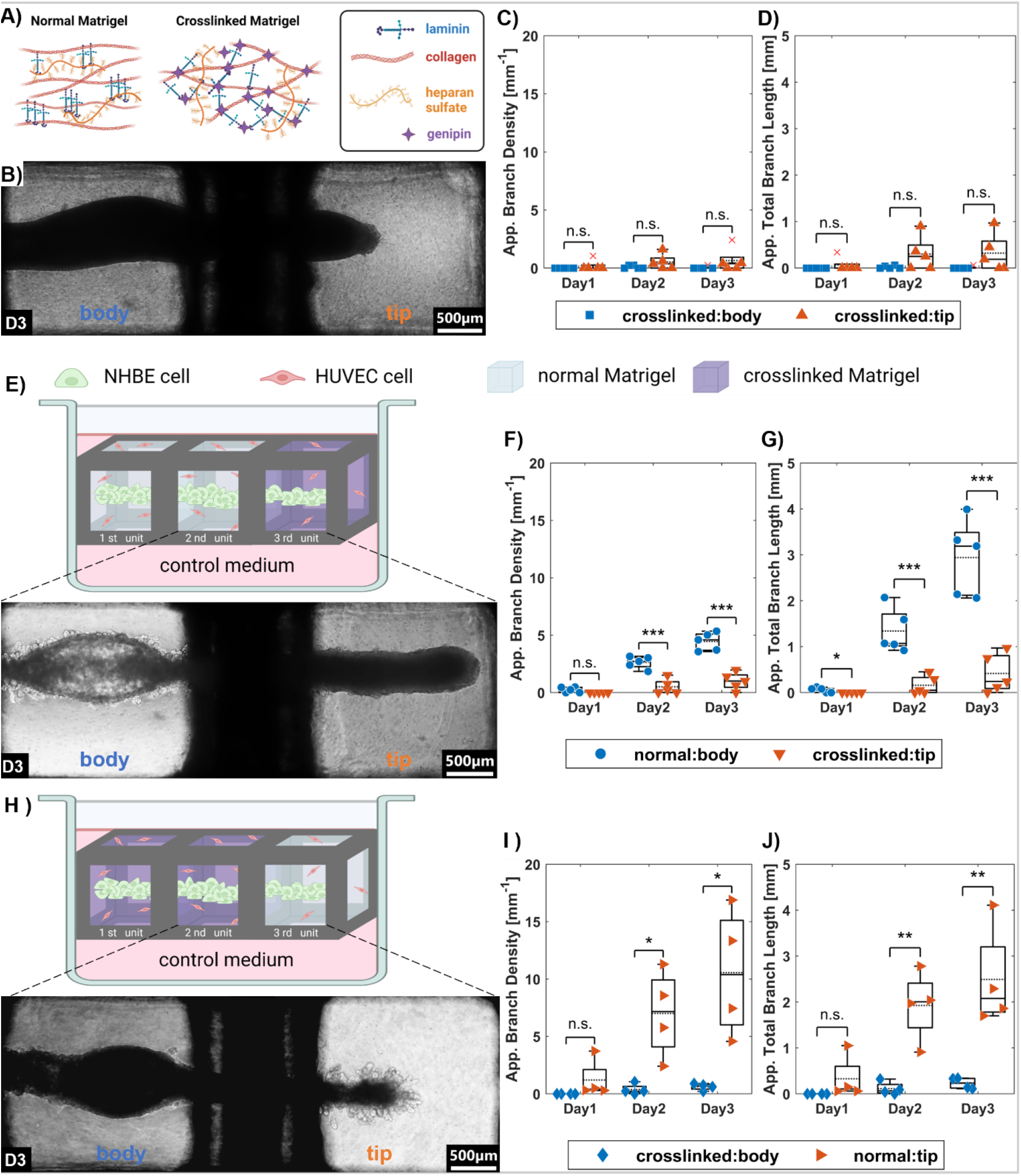
Localization of crosslinking condition and the controllability of localized branching morphogenesis. **A)** Constituent proteins in a normal Matrigel spontaneously undergo reversible physical arrangement to form a stable soft hydrogel, whereas the addition of genipin permanently crosslinks Matrigel with covalent bonds among these proteins. **B)** NHBE tubular tissues in crosslinked Matrigel substantially contracted its tip and formed no branches on the tube surface, as indicated by **C,D)** low apparent branch density and total branch length. **E,G)** rd When localizing the crosslinked Matrigel at the 3^rd^ unit, only the tip part of NHBE tissues did not branch as indicated by **I,J)** significantly lower apparent branch density and total branch length. **F,H)** Inversely, when rd localizing normal Matrigel in the 3^rd^ unit, only the tip parts branched out with **K,L)** significantly higher apparent branch density and total branch length. Data are presented in dot plot with boxplot (x: outlier, solid line: median, dash line: mean). Statistical significances are evaluated with one-way ANOVA with Tukey-Framer post-hoc test (n.s.: no significant difference, *: *p*<0.05, **: *p*<0.01, ***: *p*<0.001). Scale bar: 500 μm.

## Discussion and Conclusion

This research achieved the localization of multiple ECM-protein solutions with the developed MultiCUBE platform to spatially guide the structural development of 3D tissue models. So far, heavily engineered processes and chemical modifications necessary for fabricating multiple-hydrogel systems (also called multicomponent, multicompartment, multilayer and multiphase hydrogels)^43–45^ limits their applications in 3D tissue modeling. However, MultiCUBE greatly simplifies the process to localize multiple ECM-protein solutions without chemical modification, which allows the embedding of biological samples in many ways. Although the exact mechanism behind the solution-trapping ability still remains unclear, we speculated the interplay among the static force balance, the edge-pinning effect^46^, and other dynamic factors (*e.g.*, gradual gelation, viscoelasticity). Understanding the mechanism warrants much more investigation beyond the scope of this research. Nonetheless, the resulting simpler platform requires only manual pipetting and handling to localize multiple hydrogels which allows researchers in many fields to adopt MultiCUBE for their purposes without any need of specialized skill. Interestingly, the development of MultiCUBE unit-based scaffold to statically localize hydrogel solutions coincided with several recent and concurrent developments of periodic lattice/mesh structure to guide solution flows^46^. This convergence of different ideas toward similar design architectures showed an increasing interest in periodic hydrogel / frame composites for other applications, *e.g.*, filtration^47^ or energy storage^48^.

The capability of MultiCUBEs to spatially guide 3D tissue morphogenesis with localized ECM-protein hydrogels opens new possibilities for 3D tissue modeling. Although some multiple-hydrogel systems have been applied in 3D cell culture to guide single-cell migration and self-organization into tissues^49,50^, very few were used to control the collective cell migration and morphogenesis within a single tissue. This research succeeded to control the morphogenesis of reconstituted epithelial tissues from human primary cells in a location-specific manner with multiple-hydrogel systems. Multiple-hydrogel localization is one among many other routes to control the structural development of 3D tissue models, *e.g.*, patterned boundary constraint^51–53^ or soluble gradient^54^. We believe that our localization could complement these techniques by providing additional spatial information from localized hydrogels to further enhance morphogenesis or drive other development processes as well.

Although MultiCUBE was designed to solve the limitations in localization, a few technical constraints persist in this platform. Firstly, 3D-printing was by far the only feasible process to fabricate MultiCUBEs, which restricts the range of frame materials and print dimension. With the current inkjet printing of photopolymerizable resin, the thinnest maneuverable surface extension of L-shaped frames was 0.5 mm in thickness with 0.02 mm surface roughness. Therefore, fabricating MultiCUBE smaller than S = 1 mm or with other processes remains a challenge to be solved. Secondly, sample characterization inside the MultiCUBE platform, especially whole-gel immunofluorescent imaging, was obstructed by hydrogel thickness due to the optical trade-off between penetration depth and resolution. Also, the L-shaped frame itself blocks some part of NHBE tissues especially at the interface between two different hydrogels. Although MultiCUBE could be easily broken by a pair of tweezers for sample retrieval, better methods to characterize samples inside the platform remain to be explored. Nonetheless, these caveats do not compromise the MultiCUBE performance in localizing multiple hydrogel solutions, but rather stimulate the topic interest which could benefit researchers from additive manufacturing, optical imaging and other fields as well.

In conclusion, multiple-hydrogel localization with the MultiCUBE platform successfully guided the location-specific structural development of 3D tissue models. This control of structural development would be a great tool to study the balance between cell-cell and cell-matrix interaction, the crosstalk among different tissue parts, and other complex processes in developmental biology. Potentially, this platform-enabled localization of multiple hydrogels could be used to control cellular differentiation in a location-specific manner as well. It opens the possibility of recapitulating the true organotypic structures and functional cellular populations for disease diagnostics and drug development in the future.

## Supporting Information

Supporting information is available from the Wiley Online Library or from the authors.

## Acknowledgement

This research was supported by funds from JSPS KAKENHI Grant numbers JP21H01299. Some illustrations were created with BioRender.com

