## Supporting Information for "Localization of multiple hydrogels with MultiCUBE platform spatially guides 3D tissue morphogenesis *in vitro*"

[1] Human Biomimetic System RIKEN Hakubi Research Team, RIKEN Cluster for Pioneering Research (CPR), Wako, Saitama, Japan.

---

### Supplementary Figure List

**Fig.S1:**
Disagreement between the value trend of $6(S^2\text{-}W^2) / S^3$ ratio and the solution-trapping ability results.

**Fig.S2:**
Success in localizing hydrogel localizations in 3x1 MultiCUBE along different orientations and filling directions.

**Fig.S3**
Solution trapping process in different MultiCUBE orientation and filling direction.

**Fig.S4:**
The effect of hydrogel compositions on tumor/angiogenesis models.

**Fig.S5:**
Details in the templating method.

**Fig.S6:**
Quantification of branching development from base-view images.

**Fig.S7:**
Comparison between the culture conditions without localization and with HBEGF(+)-Matrigel localization.

**Fig.S8:**
Comparison between the culture conditions with and without crosslinked-Matrigel localization.

**Fig.S9:**
Cross-comparison of NHBE branching in the same hydrogel condition at different units.

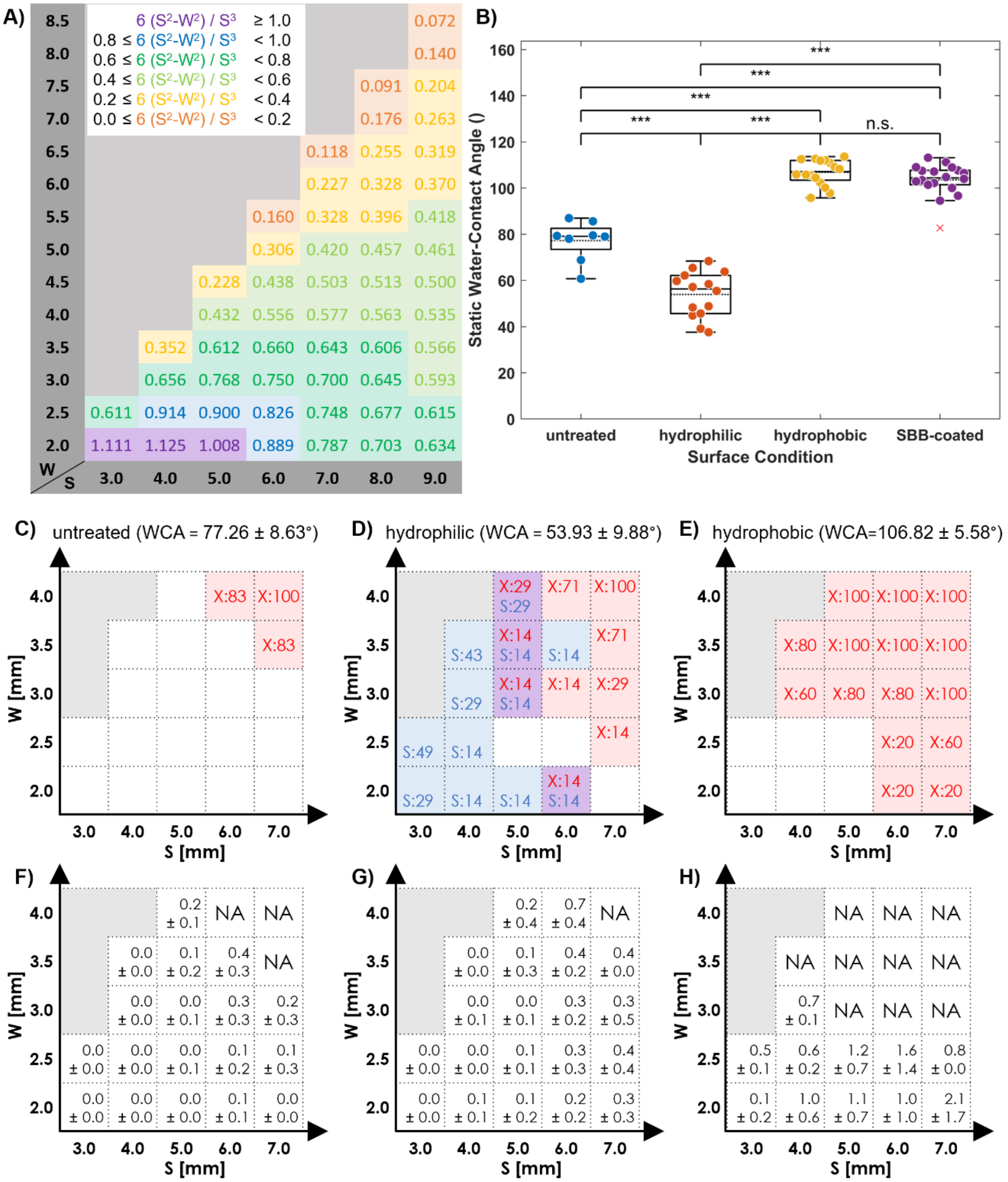


**Fig.S1: Disagreement between the value trend of $6(S^2\text{-}W^2) / S^3$ ratio and the solution-trapping ability results.** **A)** The value trend of area-to-volume ratio $6(S^2\text{-}W^2) / S^3$. **B)** The surface wettability was determined by static water-contact angle (WCA) measurement, which showed that SBB-coated MultiCUBE surface were hydrophobic. WCA measurement of untreated, hydrophilic, hydrophobic, and SBB-coated units was calculated from n = 8, 14, 16, and 18, respectively. Data are presented in dot plot with boxplot (x: outlier, solid line: median, dash line: mean). Statistical significances are evaluated with one-way ANOVA with Tukey-Framer post-hoc test (n.s.: no significant difference, *: $p<0.05$, **: $p<0.01$, ***: $p<0.001$). **C,D,E)** The occurrence percentage of spreading (S) and dripping (X) across different (S,W) grid at different surface wettability. **F,G.H)** The mean ± standard deviation of hanging-base height across different (S,W) grid at different surface wettability. Note that hydrophobic units whose dripping percentage were X:80 in **E)** corresponded to NA in **H)** because hydrogel solutions developed a hanging base only once out of five replicates (thus 20% hanging base and 80% spreading).

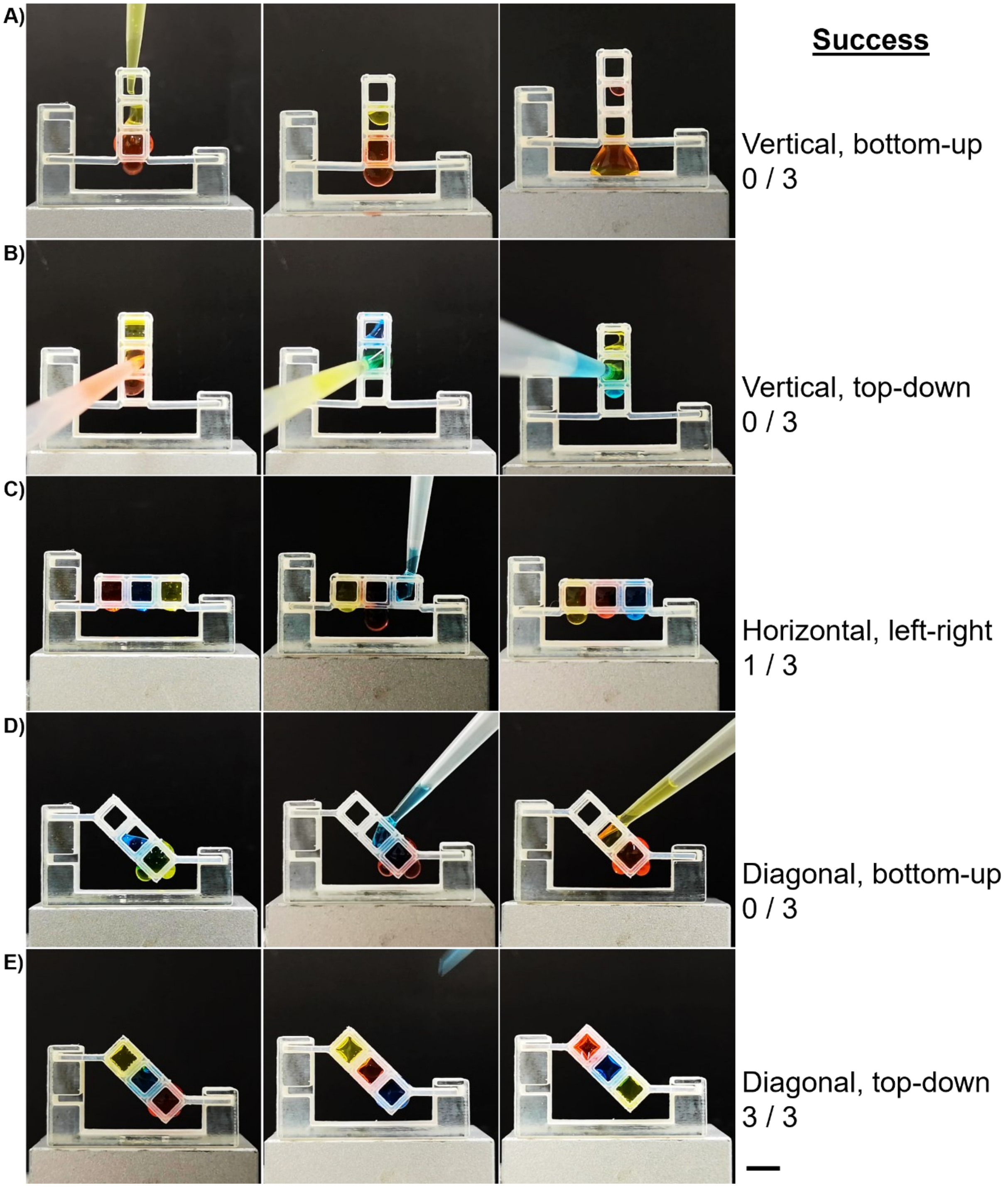


**Fig.S2: Success in localizing hydrogel localizations in 3x1 MultiCUBE along different orientations and filling directions.**

**A)** Vertical orientation, bottom-up direction; **B)** vertical orientation, top-down direction; **C)** horizontal orientation, left-right direction; and **D)** diagonal orientation, bottom-up direction all led to failure to localize hydrogel solutions. **E)** Diagonal orientation, top-down direction led to successful localization of three hydrogel solutions. Note that the convex base conformation of hydrogel solutions still appeared at the bottommost unit. For each unit, S = 4 mm and W = 3 mm. Scale bar: 5 mm.

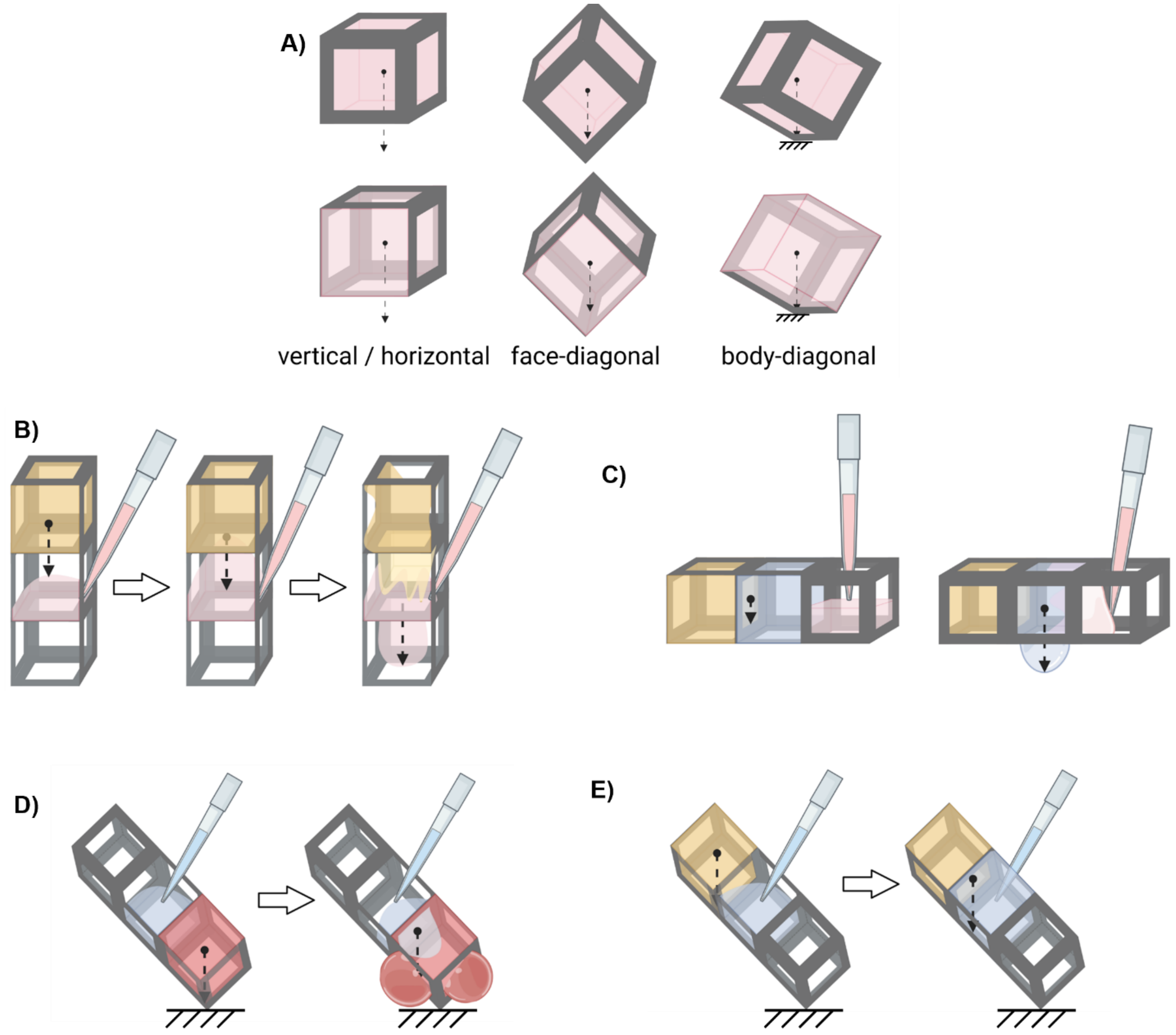


**Fig.S3: Solution trapping process in different MultiCUBE orientation and filling direction.**
**A)** Either vertical or horizontal orientation directs CG vector (dashed arrow) of trapped solutions in each MultiCUBE unit through the open base, whereas the face-diagonal and body diagonal orientation directs the vector toward the edge and the corner of MultiCUBE unit, respectively. **B)** Vertical orientation always fails in trapping solution because of CG vector shifting without any physical barrier, as well as the solution tendency to cohere before complete surface adhesion. **C)** Horizontal orientation usually fails in trapping when the third solution is being filled because the liquid cohesion before complete surface adhesion shifted CG vector to the open base on the 2$^{nd}$ unit. **D)** Although the CG in the body-diagonal orientation always direct toward the corner (and the edge during CG shifting), a strong cohesion between the 2$^{nd}$ solution and the 1$^{st}$ solution always led to solution shifting downward when filling from bottom up. **E)** However, solution filling from top down in the body-diagonal orientation allowed the 2$^{nd}$ unit to establish most of the surface-liquid adhesion before liquid-liquid cohesion could be established, thus leading to a successful localization. Some frame parts were removed to facilitate viewing.

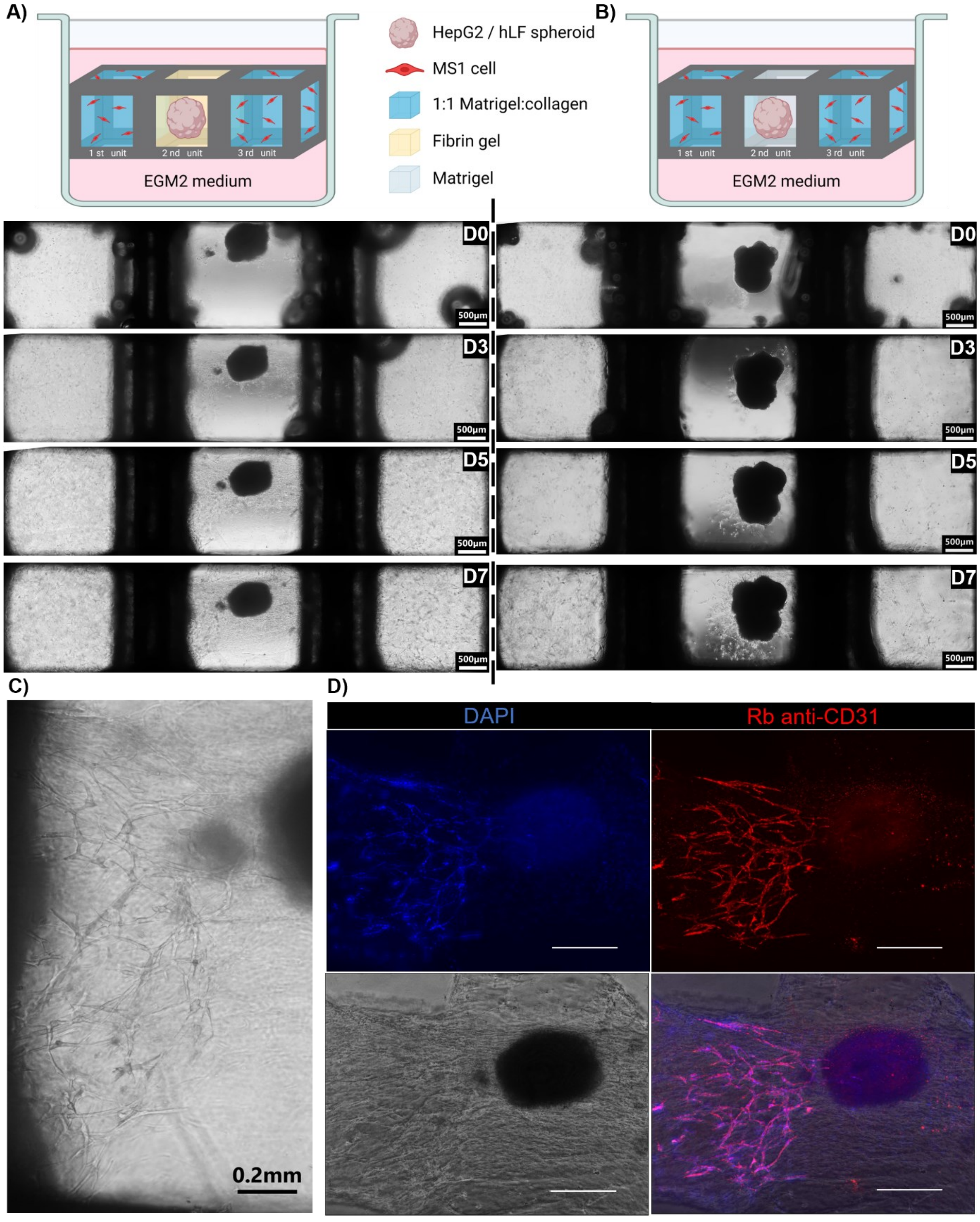


**Fig.S4: The effect of hydrogel compositions on tumor/angiogenesis models.**

**A)** Culturing HepG2/hLF tumor spheroid in fibrin gel allowed MS1 from the 1st and 3rd units to migrate toward the spheroid within seven days of culture. **B)** Culturing the tumor spheroid in Matrigel prevented the single-cell migration of MS1. Note that hLF began to spread outward from the inside of the spheroid instead. **C)** The capillary-like structure was visible under 10x bright-field microscopy. **D)** Immunofluorescent staining confirmed the capillary-like structure near the tumor spheroid in fibrin gel originated from the self-organization of MS1 cells. Scale bar: 500 µm.

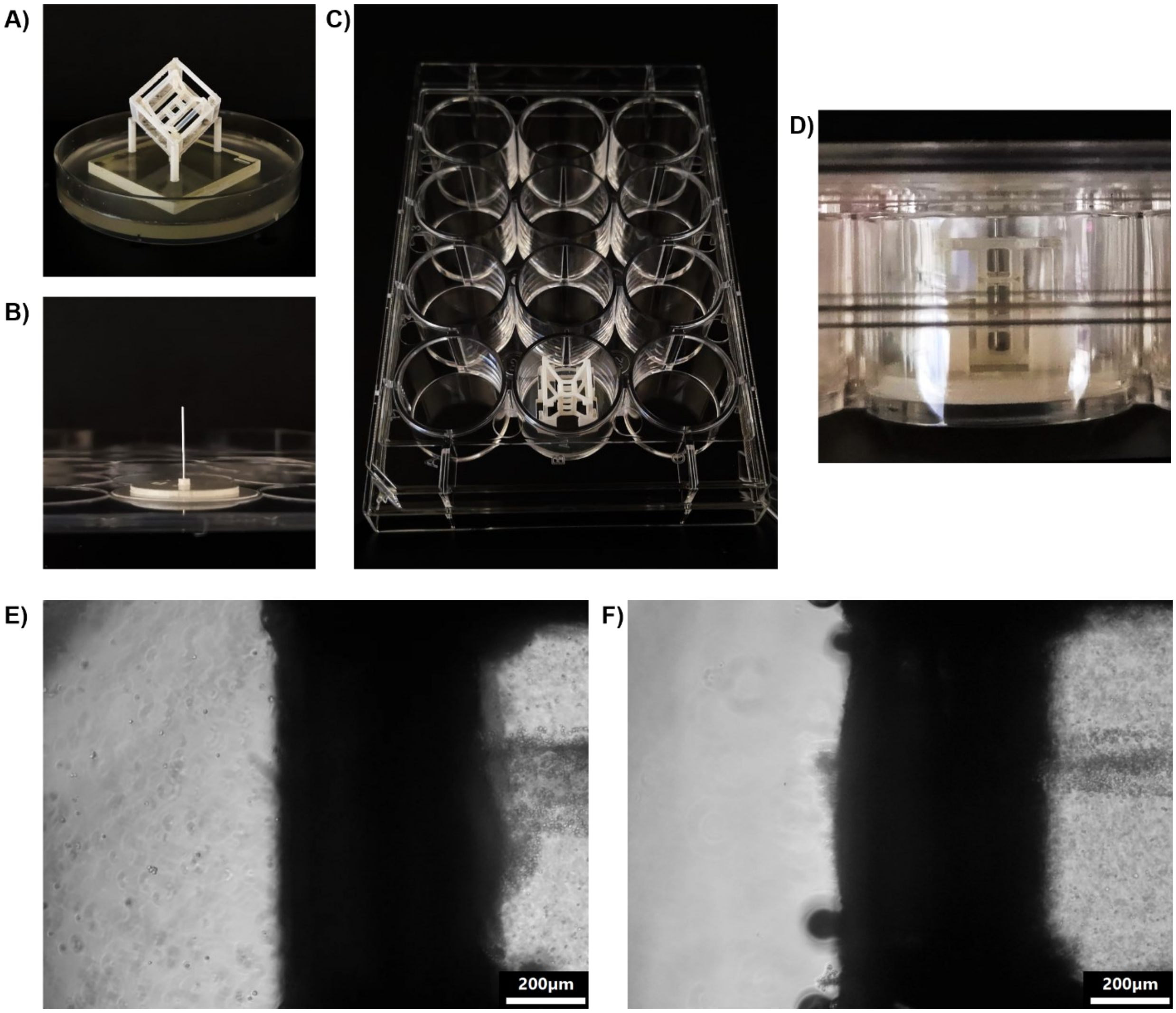


**Fig.S5: Details in the templating method.**
**A)** MultiCUBE was first placed in a body-diagonal holder and hydrogel solutions were filled into each unit. **B)** A straight aluminum wire of 0.3 mm diameter was fixed to a 3D-printed jig attached on the lid of 12-well plate such that the wire aligned through the center of the well. **C)** The filled MultiCUBE was placed vertically into a jig inside a 12-well plate such that all three units aligned to the center of the well. **D)** The lid was carefully placed to close the well plate and to insert the wire through the center of all units. **E)** Without the inlet-sealing step, cells leaked out of the molded space into the medium upon submersion. Not only the density of the tissue was disrupted, but the leaking cells also eventually grew on the well surface and consumed the nutrients and growth factors from the medium. **F)** Sealing the inlet with ~2.5 µl drop of hydrogel solutions (corresponding to the solution in the $1^{st}$ unit) sufficiently prevented the leakage of molded cells but also pushed molded cell from the $1^{st}$ unit toward the $2^{nd}$ unit.

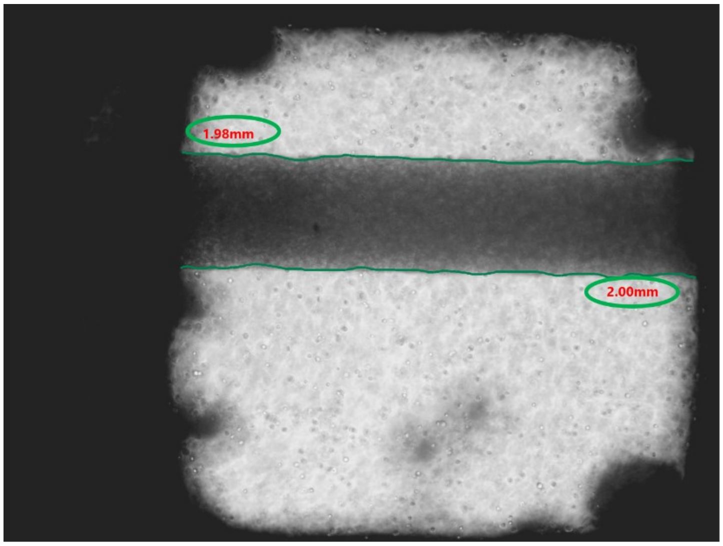


Apparent Tissue Tube Perimeter (P)
= 1.98 + 2.00
= 3.98 mm

Apparent Branch Number (N)
= 0

Apparent Total Branch Length ($\sum_{i=1}^{N} L$)
= 0

Apparent Branch Density (N / P)
= 0

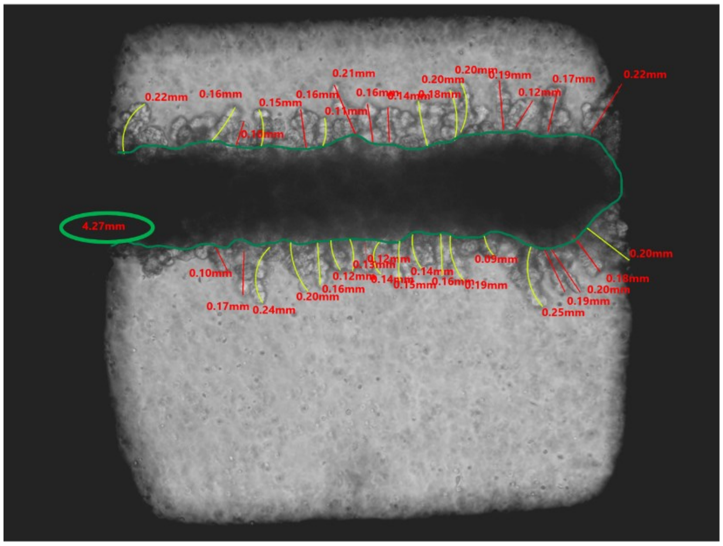


Apparent Tissue Tube Perimeter (P)
= 4.27 mm

Apparent Branch Number (N)
= 36

Apparent Total Branch Length ($\sum_{i=1}^{N} L$)
= 10.09 mm

Apparent Branch Density (N / P)
= 36 / 4.27
= 8.43 $mm^{-1}$

**Fig.S6: Quantification of branching development from base-view images.**
Images were analyzed with AdvanVision software where the apparent outline of NHBE tube (green) was measured as the apparent tissue tube parameter. The apparent branch length was approximated from the tip of each branch to the outline of NHBE tube (yellow green, red).

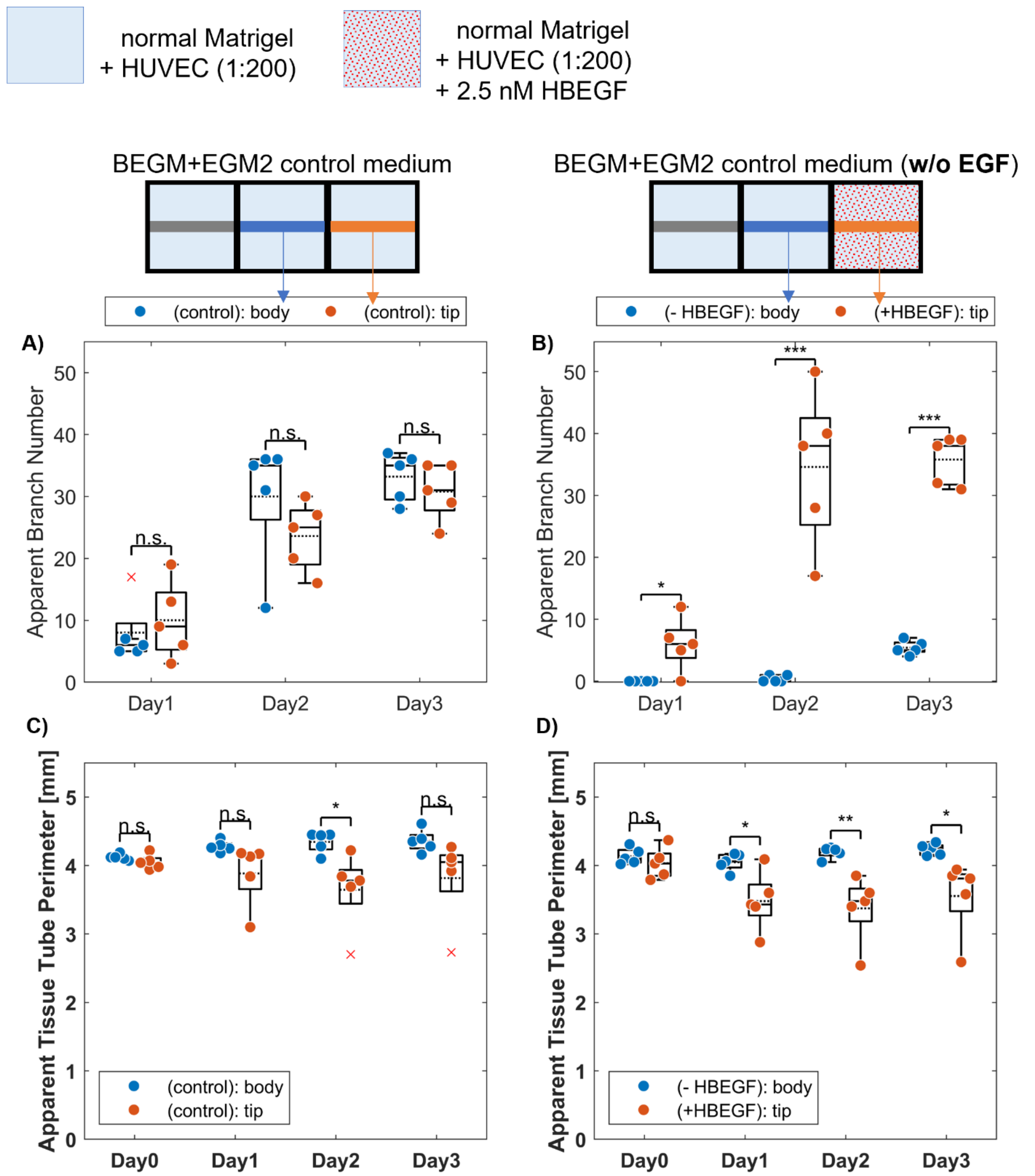


**Fig.S7: Comparison between the culture conditions without localization and with HBEGF(+)-Matrigel localization.**

**A,B)** The condition without localization showed no significant difference in the apparent branch number between the body and the tip parts, whereas the tip parts in the localizing condition showed significantly higher number than the body part in all three days of culture. **C,D)** The apparent perimeter of the tip part in both conditions gradually decreased over three days of culture due to the tip contraction toward the 2nd unit. Data are presented in dot plot with boxplot (x: outlier, solid line: median, dash line: mean). Statistical significance are evaluated with one-way ANOVA with Tukey-Framer post-hoc test (n.s.: no significant difference, *: $p<0.05$, **: $p<0.01$, ***: $p<0.001$).

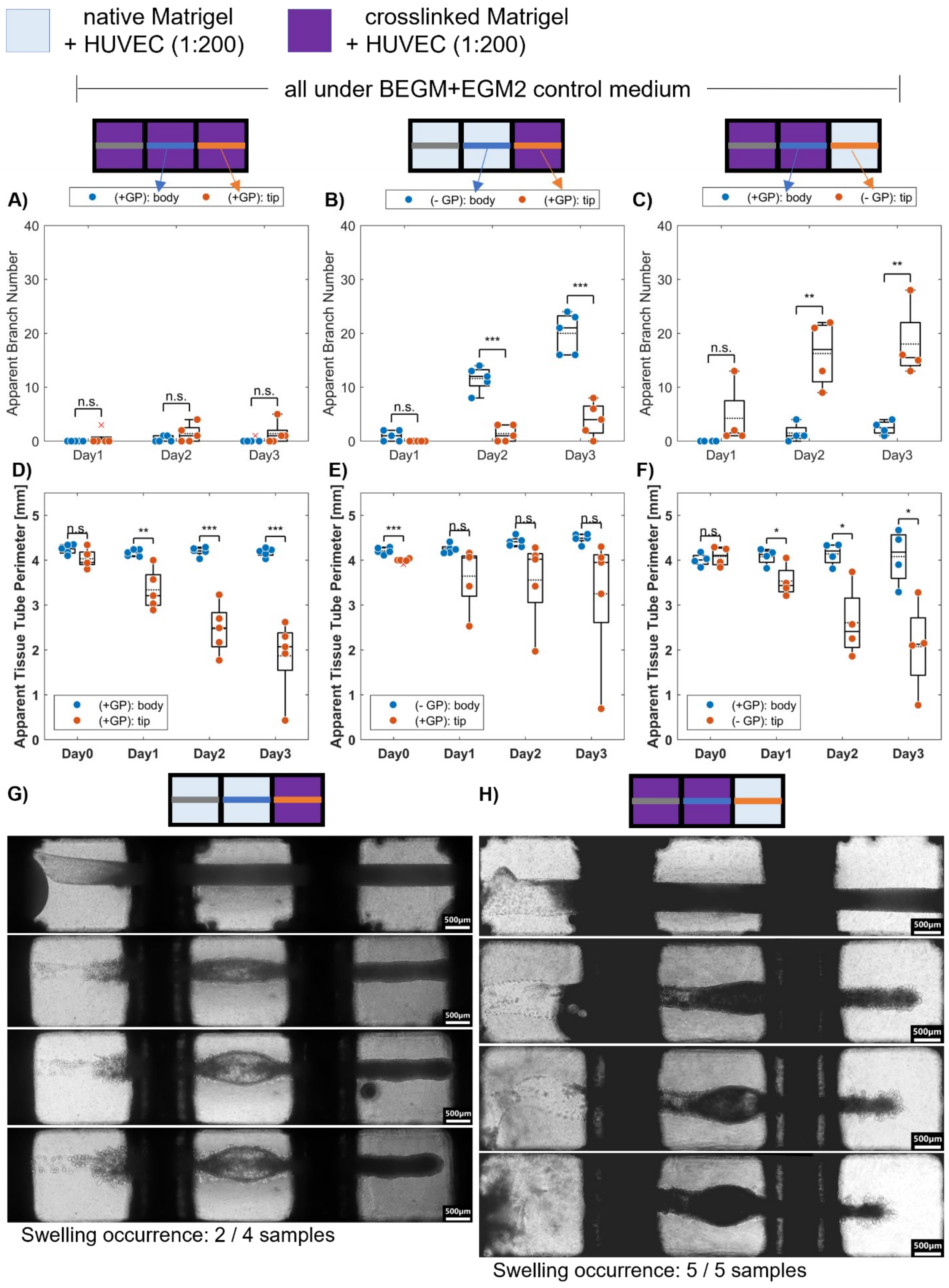
native Matrigel + HUVEC (1:200)
crosslinked Matrigel + HUVEC (1:200)
all under BEGM+EGM2 control medium
A)
(+GP): body
(+GP): tip
B)
(- GP): body
(+GP): tip
C)
(+GP): body
(- GP): tip
Apparent Branch Number
Day1
Day2
Day3
D)
E)
F)
Apparent Tissue Tube Perimeter [mm]
Day0
Day1
Day2
Day3
G)
H)
500μm
Swelling occurrence: 2 / 4 samples
Swelling occurrence: 5 / 5 samples

**Fig.S8: Comparison between the culture conditions with and without crosslinked-Matrigel localization.**
**A,B,C)** The culture conditions in crosslinked-Matrigel without localization resulted in low apparent branch number in both body and tip parts with no significant difference, whereas the conditions with crosslinked-Matrigel localization showed significant higher apparent branch number in normal Matrigel than in crosslinked Matrigel. **D,E,F)** The apparent tissue tube perimeter showed that the tip in **E)** the localizing condition contracted less than the tip in **F)** the inverted-localizing condition. **G,H**) In both localizing and inverted-localizing configuration of crosslinked Matrigel, the body parts of NHBE tissue swelled gradually over three days. Data are presented in dot plot with boxplot (x: outlier, solid line: median, dash line: mean). Statistical significance are evaluated with one-way ANOVA with Tukey-Framer post-hoc test (n.s.: no significant difference, *: $p<0.05$, **: $p<0.01$, ***: $p<0.001$).

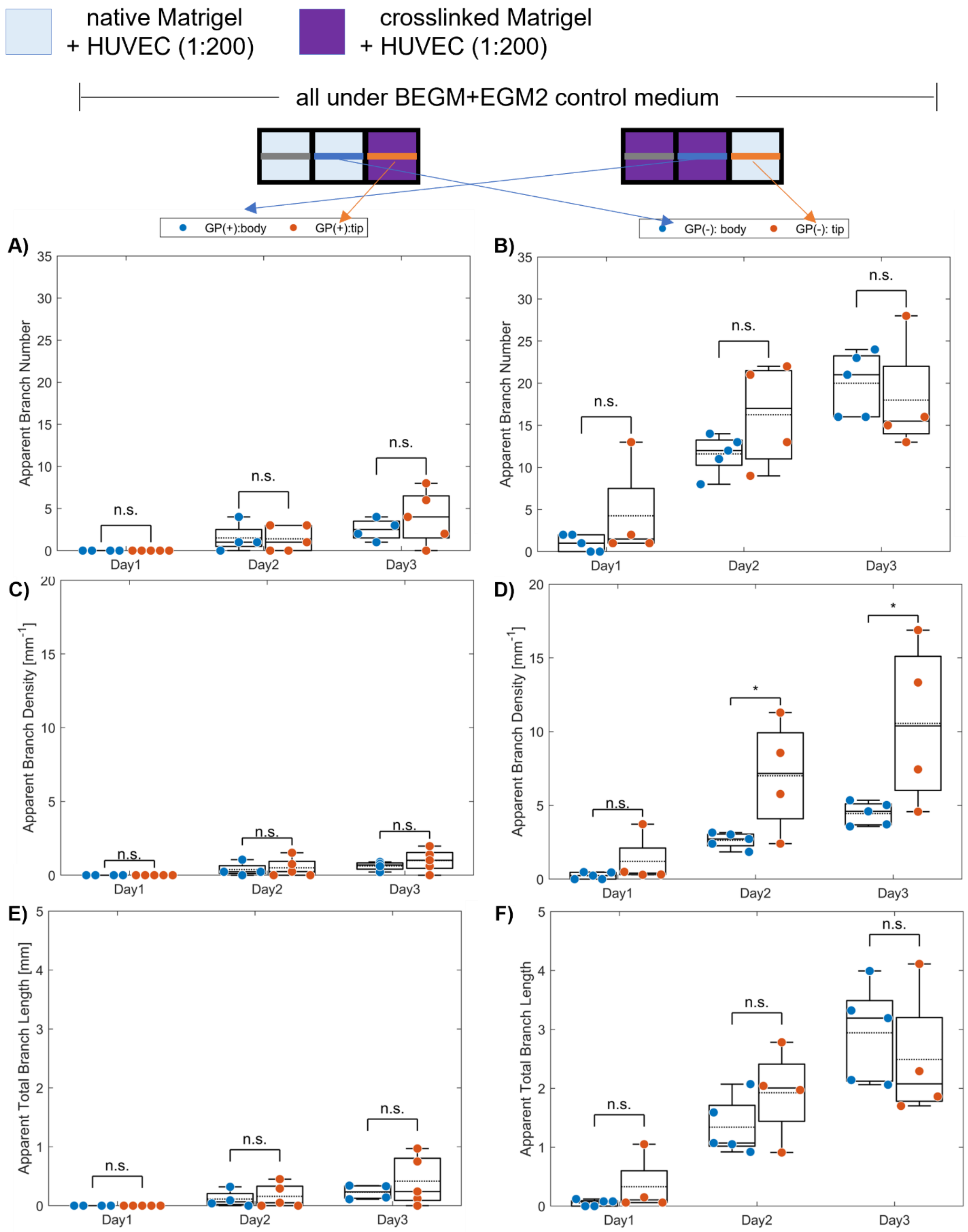

native Matrigel + HUVEC (1:200)
crosslinked Matrigel + HUVEC (1:200)
all under BEGM+EGM2 control medium
GP(+):body
GP(+):tip
GP(-): body
GP(-): tip
A)
Apparent Branch Number
B)
Apparent Branch Number
C)
Apparent Branch Density [mm$^{-1}$]
D)
Apparent Branch Density [mm$^{-1}$]
E)
Apparent Total Branch Length [mm]
F)
Apparent Total Branch Length
Day1
Day2
Day3
n.s.
*

**Fig.S9: Cross-comparison of NHBE branching in the same hydrogel condition at different units.**
**A)** The apparent branch number, **C)** branch density and **E)** total branch length of different tissue parts in crosslinked hydrogels showed no significant difference over three days of culture. **B)** The apparent branch number and **F)** total branch length of different tissue parts in normal hydrogels showed no significant difference over three days of culture. Only **D)** the apparent branch density in normal hydrogels showed significant difference on Day 2 and Day 3 due to more contraction of the tip part toward the $2^{nd}$ unit. Data are presented in dot plot with boxplot (x: outlier, solid line: median, dash line: mean). Statistical significance are evaluated with one-way ANOVA with Tukey-Framer post-hoc test (n.s.: no significant difference, *: $p<0.05$, **: $p<0.01$, ***: $p<0.001$).
